# Effects of Social Status on Reproductive Success, Physiology and Maternal Care in Socially housed Female Rhesus Macaques

**DOI:** 10.1101/2025.10.31.685853

**Authors:** K. Bailey, T. Jonesteller, Z.A. Kovacs-Balint, J. Bachevalier, M.C. Alvarado, M.E. Wilson, J. Raper, M.M. Sanchez

## Abstract

This study examined the long-term effects of low social status on reproductive success and seasonal changes in reproductive and stress hormones of adult female rhesus macaques. Rhesus are a matriarchal, matrilineal species that form hierarchical social hierarchies maintained by aggression. Thus, animals in the bottom of the hierarchy (low social status: subordinate) experience high levels of aggression from those in higher social ranks and therefore must remain vigilant. Social subordination generates chronic psychosocial stress and could negatively impact reproductive function. Twenty-seven adult female rhesus monkeys (13 dominant -DOM-, 14 subordinate -SUB-; 8-11 years old) were studied for stress neuroendocrine function -measuring basal plasma levels of cortisol-, as well as for reproductive endocrine function -measuring basal levels of estradiol (E2). In addition, we examined the reproductive success rates of these females (defined as number of live births/years with male access), their success rates raising infants that survived to 1 year, and the characteristics and effectiveness of their maternal care. Our findings show significantly lower reproductive success rates in SUB than DOM females, blunted E2 seasonal changes, driven by higher E2 levels in the anovulatory season and more attentive maternal care of offspring early in life than dominant females. Age at first birth was negatively associated with infant survival rate across ranks. Interestingly, higher levels of reproductive hormones (E2) during the anovulatory season predicted lower reproductive success, although the effect was driven by animals lactating at that point. Overall, these findings suggest that subordinate female macaques show a phenotype consistent with impaired reproductive function.

**Simple Summary:** This study examined the long-term effects of low social status on reproductive success and seasonality of reproductive and stress physiology of female rhesus monkeys. The despotic rhesus social hierarchy is matrilineal and maintained by aggression. Males leave their natal group in bachelor groups around puberty, but females remain, so social status of group members is passed down the female line. Rhesus with low social status (subordinate) experience significantly more aggression and harassment than their higher-ranking counter parts (dominant). This makes rhesus macaques an ideal model for studying chronic psychosocial stress and its long-term effects. Our findings show significantly lower reproductive success rates (number of live births based on access to males; lower infant survival rates) in subordinate than dominant females. Subordinate females also show blunted seasonal changes of the gonadal hormone estradiol (E2), driven by higher E2 levels in the anovulatory season than dominant animals, as well as more attentive maternal care of offspring early in life than dominant animals. Age at first birth was negatively associated with infant survival rate across ranks. Interestingly, higher levels of reproductive hormones (E2) during the anovulatory season were associated with lower reproductive success, although the effect was driven by animals lactating at that point. Overall, these findings suggest that female macaques with lower social status show a phenotype consistent with impaired reproductive function, though the biological mechanisms involved need to be further explored.

## Introduction

Social stress has an inhibitory impact on fertility and its neuroendocrine regulation, both in women [1, 2] and nonhuman primates -NHPs-[3-8]. In particular, low social status -social subordination-in female rhesus monkeys results in chronic social stress [9, 10] that affects reproductive endocrinology [9, 11-13] . Chronic stress, including chronic social stress like that experienced by those of a low socio-economic status (SES) has been linked to issues with reproductive success, obstetric problems, and less success raising healthy infants [14, 15]. Despite this strong link between low SES and issues with reproductive health and function in women, studies in humans are confounded by factors such as access to health care, income and education that are very difficult to disentangle from the question of whether low SES, and the stress associated with it, is truly the leading cause. Translational NHP models provide us with a great alternative to address these questions with more experimental control over these variables. In particular, rhesus monkeys are a great NHP model with complex social hierarchies and stratified social status that result in chronic psychosocial stress for females at the bottom of the social stratum (low social status or social subordinates -SUB-). Rhesus monkeys live in female societies with matrilineal and matriarchal social structures where infants assume their mothers’ social status and males have a separate social hierarchy [16]. The dominance hierarchies are maintained through aggression, where low status females receive more aggression and harassment from dominant animals, less affiliative behavior, and exhibit higher stress and anxiety levels than dominants [17, 18]. Due to this dominance hierarchy, the incidence of fighting diminishes, as subordinates typically exhibit submissive behaviors -particularly withdrawal- and abstain from retaliatory physical aggression, thereby acknowledging their place in the social hierarchy [19, 20]. This results in hypothalamic-pituitary-adrenocortical (HPA) axis over-activation and chronic-stress-related phenotypes in subordinates -e.g. emotional dysregulation and higher expression of genes related to inflammation-[9, 18, 21, 22]. The effects of social subordination are present early in development, with low-ranking animals starting to receive aggression during infancy, although not before weaning [23]. Thus, by the time subordinates are juveniles, they already show elevated levels of the stress hormone cortisol [24]. Additionally, immunological and hormonal signals passed in breastmilk from subordinate mothers could also contribute to early onset of alterations in infant development, including brain, physiology and behavior [25, 26]. Given that social status remains relatively constant for females throughout their life, in contrast to males who are often placed in different social groups and encounter changes in social rank, [18] the main question in this study was whether females’ social rank may have early programming effects to allow them to adapt to a life-long hostile social environment and relationships with others in the group, though resulting in cumulative exposure to stress, both of which will negatively affect their reproductive physiology and success rates to produce and raise healthy infants.

Unlike humans, rhesus macaques experience distinct reproductive seasonality, in which they ovulate and breed in the fall season and give birth in the spring [27]. This coincides with the end of the monsoon season in India and southeast Asia, where these animals originate. Each female typically undergoes two to six ovulation cycles per season [28] and for the most part, gives birth to one infant each year [29]. Her pregnancy lasts around 165 days, and she then lactates for approximately four to six months, with the gradual weaning of the infant coinciding with the start of the next breeding period [30, 31]. The female rhesus gives birth to an average of 7.1 infants throughout her life, although this number can vary greatly depending on the individual [32]. Outside of this seasonality, the rhesus macaque reproductive system, both physiologically and behaviorally, serves as an excellent model of human reproductive processes.

Chronic social stress, including that experienced by rhesus monkeys with low social status (subordinate), has been shown to affect many aspects of primate biology, including female reproduction [6, 12, 33, 34];. This occurs through various mechanisms including suppression of reproductive hormone secretion (FSH, LH, E2, P) by the hypothalamic-pituitary-gonadal(HPG) axis [35]. Despite the known link between stress and female reproductive function suppression, the specific role of elevated levels of cortisol due to activation of the hypothalamic-pituitary-adrenal (HPA) stress neuroendocrine axis is unclear, particularly for low-ranking individuals and across annual seasons. Thus, some studies have reported higher cortisol levels in low-than high-ranking rhesus females [36, 37], while others have reported more nuanced effects of stress hormones that depended on reproductive state (i.e. higher stress-induced cortisol responses in lactating than cycling rhesus females, particularly in low-ranking individuals) [38]. Thus, two main questions in this study were to examine (1) whether differences in seasonal levels of E2 related to social rank were associated with reproductive success of female rhesus monkeys and (2) associations between seasonal levels of activity of the HPA and HPG axes by measuring baseline blood levels of E2 and cortisol during the anovulatory, mating and birthing seasons. The complex social structure and strong mother-infant bonds exhibited by the rhesus macaque also make them an excellent model for studying maternal behavior. Typical maternal behavior includes frequent contact, cradling, and proximity to the infant, as well as restraining and retrieving the infant in response to threats [39]. These behaviors have been shown to vary by individual depending on factors like infant age, group size, and maternal rank, with more dominant females spending less time in contact with their infants and exhibiting less protective behaviors [40, 41].

The goal of this study was to investigate possible long-term effects of chronic social subordination on reproductive function, physiology and maternal care in socially housed female rhesus macaques at the Emory National Primate Research Center Field Station breeding colony.

## Methods

### Subjects and Housing

Twenty-seven adult female rhesus macaques (*M. mulatta*) were studied longitudinally from young adulthood through middle age (8-11 years), as part of a larger study examining the cumulative effects of social subordination stress on biological and neurocognitive accelerated aging (NIH/NIA R01 AG070704). Subjects lived in social groups at the Emory National Primate Research Center (ENPRC) Field Station, Emory University (Lawrenceville, GA). They were provided standard high fiber, low fat monkey chow (Purina Mills LLC, St Louis) twice daily, fruits and vegetables and water ad libitum. From birth through early adolescence, subjects were studied longitudinally as part of a different NIH funded project (NICHD R01 HD077623) that examined the effects of postnatal social subordination stress (ELS) and obesogenic diet on development. Subjects were born of multiparous mothers with a consistent history of competent maternal care in large social groups with multiple matrilines. These mothers ranked either in the top one-third or bottom one-third of their social groups and were thus characterized as Dominant (DOM) or Subordinate (SUB), respectively. Additionally, each animal was assigned a relative social rank number between 0 and 1 that represented her rank position divided by the total number of adult females in the social group (e.g. the alpha female was 1 out of 100 animals=0.01; adult males had a separate social hierarchy). DOM animals, therefore, have low relative ranks while SUB animals have high relative ranks. In order to disentangle the effects of postnatal social subordination stress from those of heritable and prenatal factors, a cross-fostering design was used in half of the infants in each DOM and SUB group, with random assignment at birth to either high- or low-ranking foster mothers in other social groups following published procedures [39]. For comparison, the remaining infants were raised by their biological mothers (see Table 1 for breakdown of experimental groups). Thus, the final breakdown of social Rank experimental groups was based on that of the mother that raised the infant (labeled as “Birth Social Rank”): DOM (n=13), corresponding to the top third of the social hierarchy and SUB (n=14), in the bottom third of the social hierarchy, counterbalanced by crossfostering (or not) and biological mother rank. Dams with a known history of neglect or physical abuse towards infants were excluded from the study. Infants, with a birth weight <450g, were also excluded from the study to avoid confounding effects of prematurity/low-birth weight on brain development [42].

**Table 1.** Breakdown of groups by social rank (dominant -DOM-, subordinate -SUB-), cross-fostering status (cross-fostered -C-, not cross-fostered -NC-), and diet (low calorie diet only - LCD-, high + LCD -Choice-)

| Rank Assigned At Birth |  |  | Diet |  |  |
| --- | --- | --- | --- | --- | --- |
|  |  |  | Choice | LCD |  |
| DOM | C | SUB→DOM | 4 | 3 | n=13 |
|  |  | DOM→DOM | 1 | 1 |  |
|  | NC | DOM→DOM | 3 | 3 |  |
| SUB | C | DOM→SUB | 1 | 0 | n=14 |
|  |  | SUB→SUB | 3 | 3 |  |
|  | NC | SUB→DOM | 1 | 4 |  |
|  |  |  |  |  | n=27 |

All pregnant females were fed a chow-based, LCD-only, healthy diet throughout pregnancy to avoid confounding effects of different prenatal dietary environments on our brain measures [43, 44]. Mother-infant pairs were randomly assigned at birth to diet groups: LCD-only condition (n=7 DOM, 6 SUB) or to a Choice condition with access to both the LCD and HCD diet (n=6 DOM, 8 SUB). Previous research has established that, in comparison to a HCD-only diet, the Choice condition is a superior translational model for the “Western” human diet, allowing for stress-induced, emotional feeding in animal models [45]. Subcutaneous radio-frequency identification (RFID) chips were implanted in the subjects’ wrists, allowing subjects’ access to and activation of automated feeders to deliver pellets of experimental diets and accurately record the Kcal consumption of each diet [46]. These chips were programmed to enable each subject to access either only LCD feeders -LCD condition- or both LCD and HCD feeders -Choice condition-. This system has been validated at the ENPRC demonstrating that subjects almost always consume a pellet of food once they have taken it from the feeder [46].

During adulthood, some of the larger social groups experienced social instability, resulting in relocation of several subjects to new social groups of varying sizes, either with or without the rest of their matriline. For this reason, 5 of the 27 females in this study have a different adult rank than birth rank (they changed social status during adulthood -subordinate to dominant-), and these rank switches were modeled in the statistical analyses.

All procedures were approved by the Emory University Institutional Animal Care and Use Committee (IACUC), in accordance with the Animal Welfare Act and the U.S. Department of Health and Human Services’ “Guide for the Care and Use of Laboratory Animals.” Furthermore, the ENPRC is fully accredited by AAALAC International.

### Reproductive Function

#### Training and capture procedures

Using positive reinforcement, subjects were trained to separate from the social group in the outdoor enclosure and move into an indoor room and then into a specialized transfer box. They were then placed into a cage with a squeeze mechanism and trained to present a leg for awake blood sample collection through saphenous venipuncture or for vaginal swabs to monitor menstrual cycle. Blood samples were collected within 10 min from disturbance of the group reflect baseline plasma levels of stress and other hormones, as previously demonstrated to be effective [39, 47-49].

#### Menarche Determination

Once each animal reached 17 months of age, they were visually monitored at least three times per week during the breeding season for any changes in sex swelling, coloration, or the presence of blood that indicated first menses (menarche[50]). They were also trained, to transfer to the cage with the squeeze mechanism two to three days per week for vaginal swabs, performed by carefully inserting a sterile cotton swab into the vaginal canal to confirm the presence of menstrual blood.

#### Reproductive success and infant survival rates. Infant birth weights. Obstetric problems

Reproductive data for each subject was compiled via lab and historical records and included: age of menarche (first menstruation); age at first birth; reproductive success rates (defined as number of live births/years with male access -i.e. presence of at least one adult male in the social group each breeding season) and infant survival rates (defined as the total number of infants surviving to at least one year divided by the total number of live births per subject; infants’ birth weight (collected within the first 48-96 hours of birth); obstetric problems, including those that required veterinary intervention (stillbirths, dystocia, retained placental tissue or placental detachment, uterine infection, caesarean section, problems with lactation). Not all females had access to a breeding male every year, due to ENPRC colony management strategic priorities for each social group or sometimes due to the loss of breeder males for clinical reasons or group manipulations due to social instabilities. For this reason, reproductive success rates were computed as the total number of live births divided by the number of years with breeding male access. More complete reproductive data was available for the last 4 recent years, when these females were assigned to our longitudinal study, including infant’s birth weight and maternal care behavior (see the following sections).

#### Seasonal Secretory Rhythms of Gonadal and Stress Hormones: Estradiol and cortisol

Basal levels of seasonal secretion of estradiol (E2) and cortisol (CORT) hormones were measured as indexes of basal activity levels of the hypothalamic-pituitary-gonadal (HPG) and hypothalamic-pituitary-adrenal (HPA) axes, respectively, in adulthood (Range: 6.21-7.49 years; 6.94±0.094 years). Because subjects lived outdoors, under natural light conditions that control their circadian/diurnal CORT and E2 secretory rhythms, morning (AM) blood collection time was selected based on daylight times published by the US Naval Observatory (https://aa.usno.navy.mil/data/RS_OneYear) across 3 seasons per year: anovulatory (gonadal suppression in female rhesus: July-September); mating season (October-December (mating season); birth season (March-April) [27]to assess for seasonal changes in baseline E2 and CORT levels. Basal AM blood samples were collected at sunrise as previously published for socially housed rhesus living outdoors [48]. All blood samples were collected in pre-chilled vacutainer tubes containing EDTA, immediately placed on ice and centrifuged for 15 minutes at 300 RPM and 4 °C. Plasma was aliquoted into sterile microcentrifuge tubes and stored at -80 °C until assay.

#### Plasma Estradiol (E2) and Cortisol Assays

Plasma concentrations of E2 and CORT were measured simultaneously by liquid chromatography-tandem triple quadrupole mass spectrometry (LC-MS/MS) in the Endocrine Technologies Core (ETC) at the Oregon National Primate Research Center (ONPRC) as previously described [51]. Briefly, 150 μl of plasma was mixed with 100 μl water containing 3.75 ng/ml E2-d5 and 7.5 ng/ml CORT-d4 internal standards and transferred to a 400-μl 96-well Isolute supported liquid extraction (SLE+) plate (Biotage, Uppsala, Sweden). Steroids were eluted with 3 X 600 μl dichloromethane, dried under nitrogen at 40°C and reconstituted in 50 μl 25:75 methanol:water. Samples were injected onto a Raptor Biphenyl reversed-phase column (100-mm X 2.1-mm, 2.7 μm; Restek, Bellefonte, PA) with an in-line guard column (Biphenyl 5-mm X 2.1-mm, 2.7 μm) at 40°C with gradient elution (mobile phase A: 0.15 mM NH_4_F; mobile phase B: methanol) using two Nexera LC-30AD pumps (Shimadzu Scientific, Kyoto, Japan). Heated electrospray injection with ultra-fast polarity switching and scheduled multiple reaction monitoring on a Shimadzu LCMS-8050 was used for detection of E2, E2-d5 (negative mode), CORT, and CORT-d4 (positive mode). *The Endocrine Technologies Core (ETC) at ONPRC is supported (in part) by NIH grant P51 OD011092 for operation of the ONPRC*.

### Maternal Care: First Month Postpartum

Focal behavioral observations of maternal care were performed on 18 mother-infant pairs (10 DOM, 8 SUB) assigned to the studies during the infant’s fourth week of life, using an adaptation of a rhesus monkey ethogram [52]. Four thirty-minute observations were performed on separate days by four coders, with two coders assigned to each infant. Prior to collection, inter-observer reliability was reached, with >90% agreement and Cohen’s Kappa exceeding 0.8. See Table 2 for ethogram.

**Table 2.** Behavioral Ethogram of maternal-infant behaviors collected during observations.

| Behavior | Definition |
| --- | --- |
| Proximity | Infant is within arm's reach of mother for at least 3 seconds |
| Contact | At least half of infant's body is in contact with mother for at least 3 seconds |
| 3 Meters | Infant is within 3 meters of mother for at least 3 seconds; It is outside of proximity but still easily reachable |
| Leave Beyond | Infant leaves outside of 3 meters of mother |
| Cradle | Ventro-ventral contact with mother's arms wrapped around infant |
| Carry | Mother carries infant while in motion; can be ventral, dorsal, or limb |
| Follow | Persistently trailing another animal (to or from infant) |
| Groom | Picking and spreading fur of another animal (to or from infant) |
| Genital Groom | Mother is inspecting or specifically grooming genital area of infant |
| Touch | Gently placing hand on infant not in context of grooming, aggression, or play |
| Eye Gaze | Making direct eye contact with another animal (to or from infant) |
| Lipsmack | Repeatedly opening and closing of lips while looking at other animal |
| Retrieve | Mother grabs or actively encourages infant to come into contact, usually in response to a perceived threat |
| Restrain | Mother physically prevents an infant from moving away by holding its leg or tail |
| Reject | Mother prevents infant from making contact or nursing |
| Play | Rough and tumble, chase play, or non-agonistic grappling play with mother |
| Punish | Mother stops infant from a certain behavior by physically removing it harshly or by mouthing it |
| Scream | High pitch, high intensity screech or loud chirp |
| Bizarre Behavior | Abnormal, non-repetitive behavior |
| Abuse | Mother drags, crushes, throws, steps on, or sits on infant, rough grooms, or otherwise causes injury to infant |

#### Data Analysis

Variables were summarized as mean 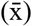 ±standard error of the mean (SEM) in figures, and effect sizes were calculated as ηp^2^. For statistical models assuming normality -all except for Spearman Rank’s correlations-, data were first checked for normality using the Shapiro-Wilk test and, if failed, data were log10-transformed and rechecked for normality. If log10-transformation did not improve the normality, the non-transformed data was utilized in the statistical model. Analyses of variance (ANOVAs) were performed to examine effects of categorical birth rank (DOM, SUB) on reproductive success, infant survival rates, and infant birth weights. Repeated Measures -RM-ANOVA were performed to examine the effects of birth rank (DOM, SUB) and seasonality (RM: anovulatory, mating, birth seasons) on E2 and CORT plasma levels. The following variables were then added as covariates in analyses of covariance (ANCOVAs) or RM ANCOVAs to control for their confounding effects: diet, adult rank, age at menarche and age at first birth; for E2 and CORT RM ANCOVAs models, additional covariates were added (infant during anovulatory season, pregnancy and trimester during birth season) to control for their confounding effects on HPG and HPA axis. Effects of birth rank on clinical reproductive and obstetric issues were not examined statistically due to their low occurrence, but the % of animals showing them in each group is presented in the Results.

Maternal care data were converted into rates per hour for frequency behaviors and minutes per hour for duration behaviors. The data were then examined to identify and exclude low occurrence behaviors (defined as ≥ 50% of the animals not exhibiting the behavior) from the statistical analyses; the following variables were excluded from the analyses: carry dorsal, carry limb, genital groom, play, follow, punish, fear grimace, infant abuse, scream, and bizarre behavior. Given the small sample size available for the remaining behavioral variables (only 18 out of 27 females) data were analyzed for associations with dam’s relative birth rank (instead of using categorical rank groups), reproductive success rate, age at first birth, and infant survival rate using bivariate Spearman’s Rank correlations because behavioral data were skewed/not normally distributed. Maternal behavior with female and male infants was analyzed separately since previous research has shown maternal care differences dependent on infant sex [39]. All statistical analyses were performed using SPSS 29.0 software, with p value set at <0.05 for significant effects.

## Results

### Reproductive Function

Shapiro-Wilk tests showed normal distributions of reproductive success rate (p=0.538) and infant birth weights (p=0.121). Neither infant survival rate nor log10-transformed infant survival rate were normally distributed (p=<0.001). E2 (p=0.014) and CORT (p<0.001) values were not normally distributed so values were log10-transformed for analysis (p=0.054).

#### Menarche onset and age at first birth

ANOVA results showed no significant effects of birth rank on age of menarche (F_(1, 19)_=0.335, p=0.570, η_p_^2^= 0.018) nor on age at first birth (F_(1, 26)_=0.578, p=0.454, η_p_^2^= 0.023). The addition of diet as a covariate did not have a significant effect on these measures or change the results. Menarche onset age was not significantly correlated with infant survival rate (r=0.304, p=0.193) or infant birth weights (r=-0.485, p=0.079), either. Age at first birth, though, was negatively associated with infant survival rate across social ranks (r=-0.387, p=0.046) but not with infant birth weights (r=-0.209, p=0.376).

#### Reproductive success and infant survival rates. Infant birth weights. Obstetric problems

ANOVA results showed a significant main effect of birth rank on reproductive success (F_(1, 26)_=4.432, p=0.045, η_p_^2^= 0.151) with lower ranking females (SUB) having lower reproductive success rates than DOM animals (Figure 1). Although neither diet nor menarche age had a significant impact on reproductive success when added separately as covariates, when they were added together as covariates to the statistical model, the effect size of birth rank on reproductive success rate increased (F_(3, 19)_=2.274, p=0.026, η_p_^2^= 0.299). None of the other covariates had a significant effect.

**Figure 1.**
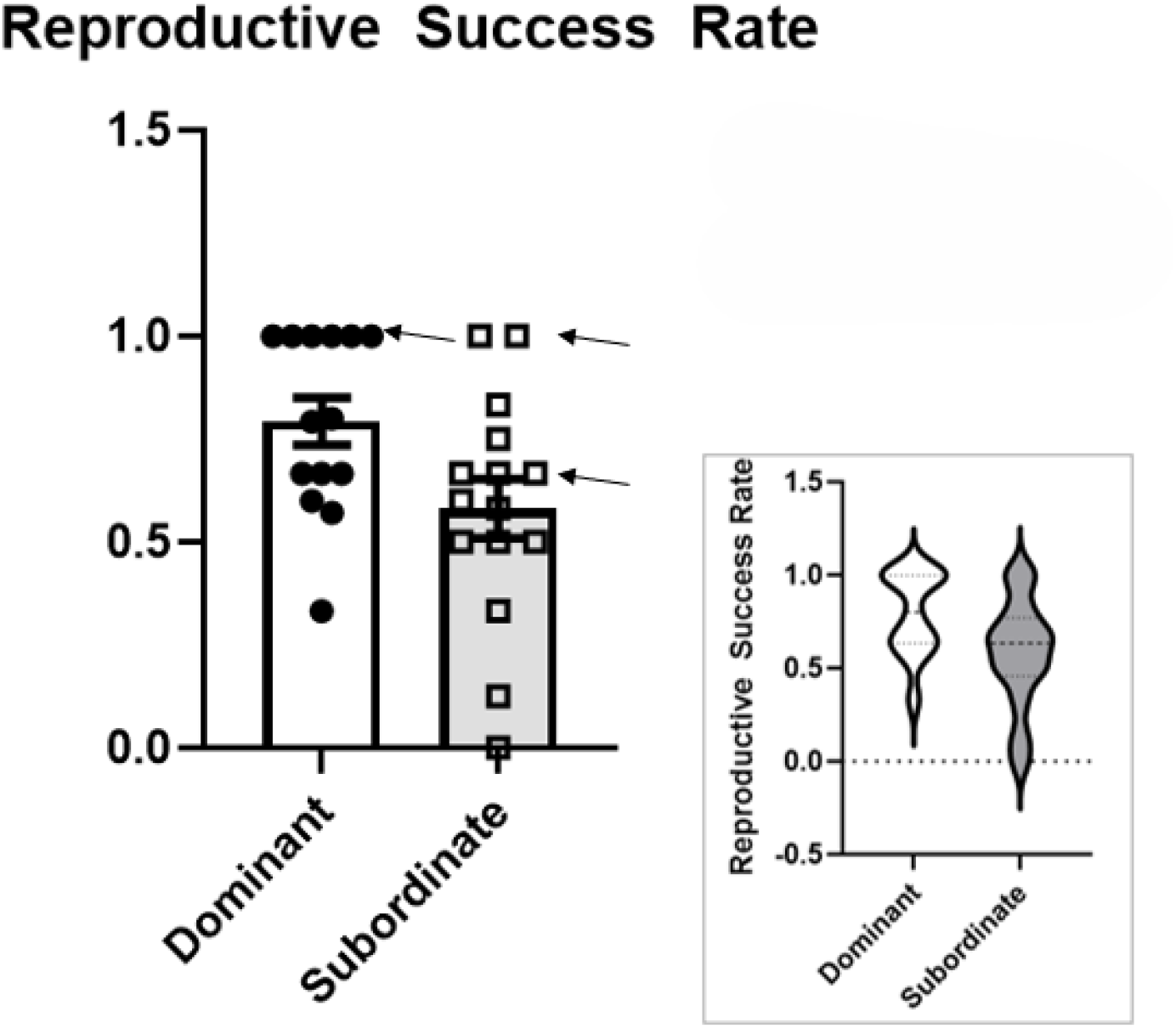
Reproductive Success Rate by birth rank (dominant -DOM-, sub -SUB-). Arrows indicate individuals whose adult rank differs from their birth rank. A significant main effect of birth rank was detected on reproductive success (F_(1, 26)_=4.432, p=0.045, η_p_^2^= 0.151) with SUB females showing lower reproductive success rates than DOMs.

Birth rank did not have a significant effect on infant survival rate (F_(1, 26)_=2.678, p=0.114, η_p_^2^= 0.097), and adding covariates to the model did not have a significant effect on the results.

ANOVA results showed no significant effects of birth rank on infant birth weights for female infants (F_(1, 16)_=0.224, p=0.643, η_p_^2^= 0.15) or male infants (F_(1, 11)_=0.083, p=0.779, η_p_^2^= 0.008), either (DOM: 490 grams ± 0.028, SUB: 471 grams ± 0.015).

Although the study was not powered to run statistical analyses for differences in obstetric problems across groups, SUB animals showed higher rates of stillbirths and lactation issues than DOM, but both groups experienced nearly equal rates of dystocia (Figure 2: descriptive data).

**Figure 2.**
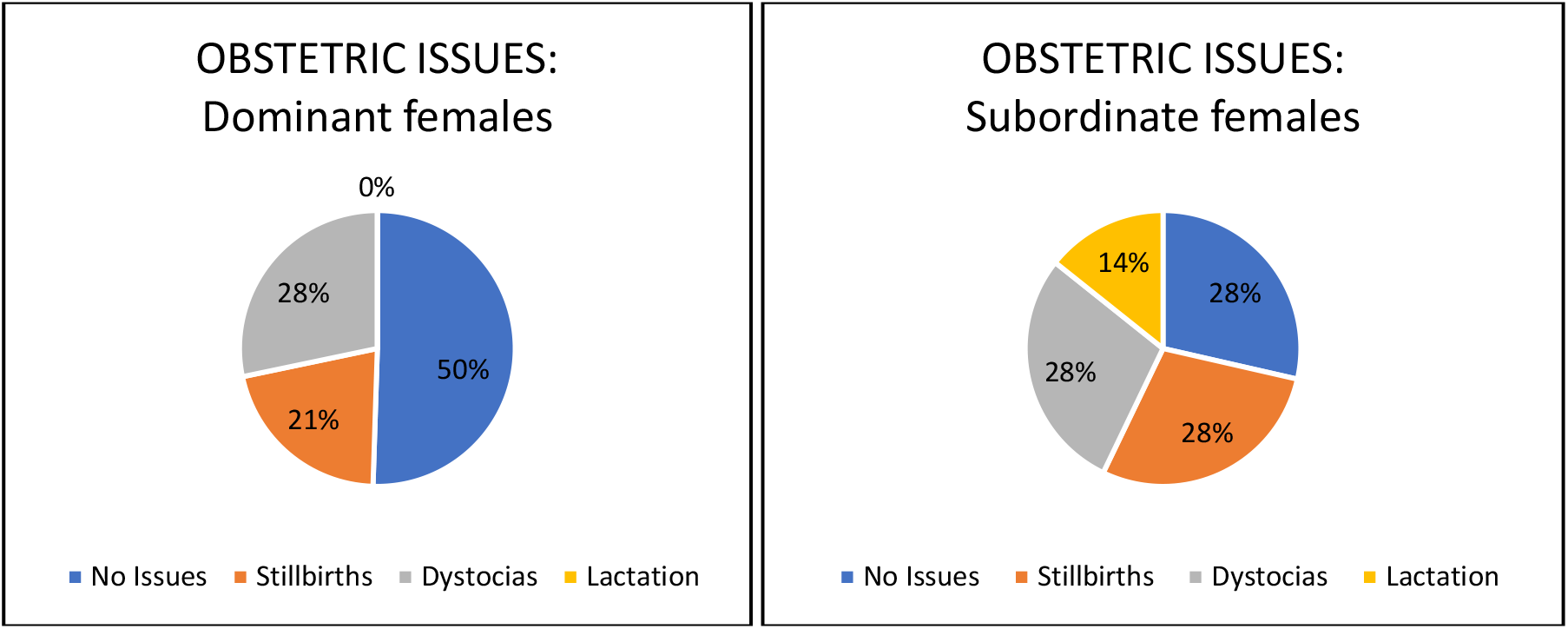
Percent of animals showing clinical obstetric issues by type based on birth rank. Subordinate (SUB) females had more obstetric and lactation issues than dominant (DOM) females.

#### Seasonal secretory rhythms of Gonadal and Stress hormones (estradiol and cortisol)

A repeated measures ANOVA showed a significant main seasonality effect (F_(1, 24)_=65.513, p=<0.001, η_p_^2^= 0.732), although with high individual variability (Figure 3A) as well as a trend toward significance of the rank x seasonality interaction effect on E2 levels (F_(1, 24)_=3.119, p=0.053, η_p_^2^= 0.115). Adding diet, adult rank, age at menarche or age at first birth as covariates to the model did not have a significant effect on the results. However, when raising an infant during anovulatory season was added as a covariate, a significant birth rank x seasonality effect emerged on E2 (F_(1, 23)_=3.525, p=0.038, η_p_^2^= 0.133), likely by controlling for the confounding effects of lactation. Although pregnancy during birth season and pregnancy trimester had significant covariate effects on E2 levels (pregnancy: F_(1, 23)_=16.591, p=<0.001, η_p_^2^= 0.419; trimester: F_(1, 23)_=14.48, p=<0.001, η_p_^2^= 0.386), no significant effect of rank emerged. When change in E2 between anovulatory and birth seasons was calculated, there was a significant rank effect (F_(1, 26)_=4.903, p=0.036, η_p_^2^= 0.164), with SUB animals showing blunted E2 increases than DOM animals. Interestingly, there was a significant negative correlation between E2 levels during the anovulatory season and reproductive success rates (r= -0.48, p=0.01), although the effect disappeared after controlling for lactation (r=-0.0306, p=0.128) and no significant association was detected between the seasonal increase in E2 and reproductive success rate (r=0.27, p=0.173).

**Figure 3.**
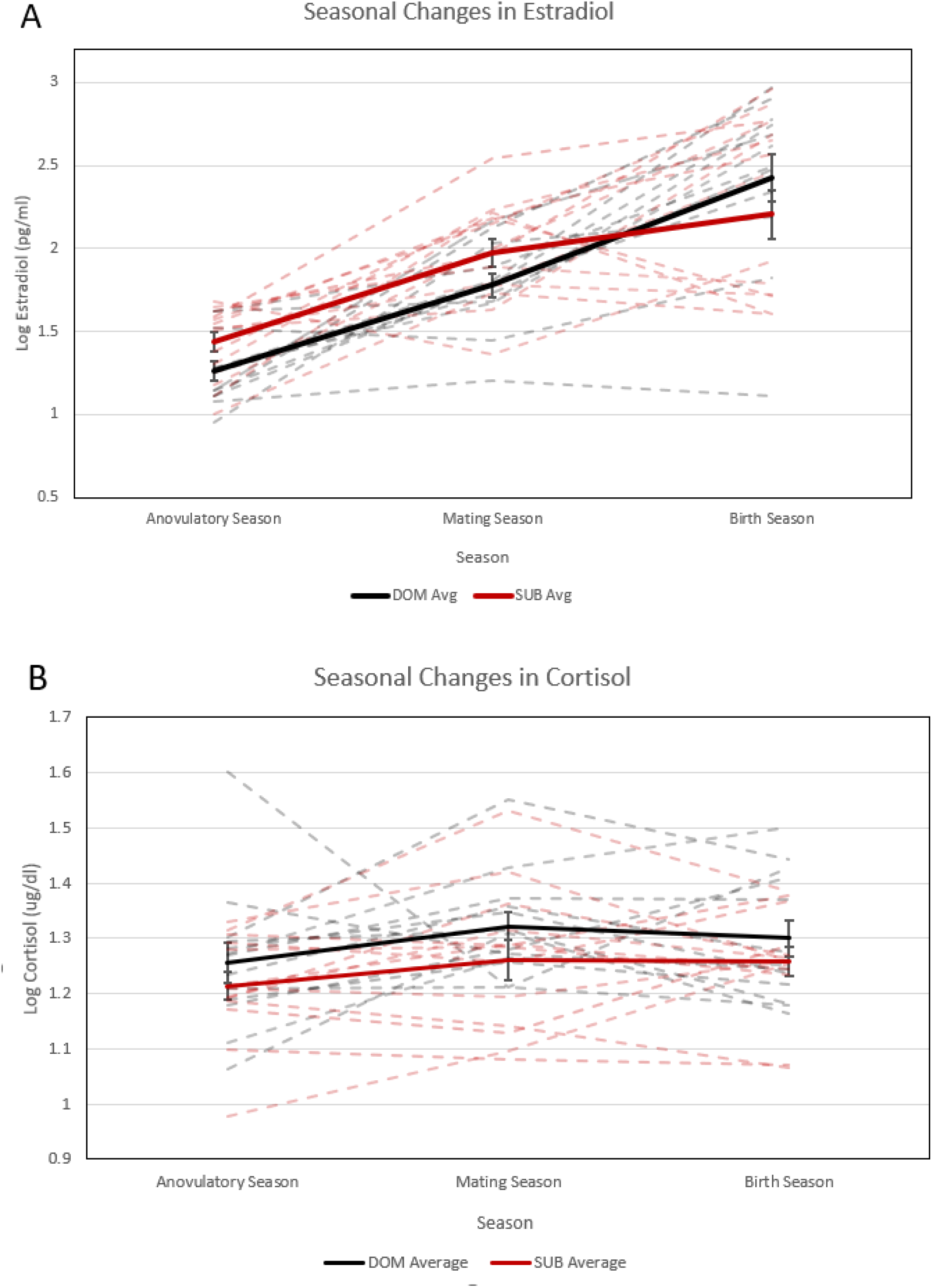
Seasonal concentrations of estradiol -E2- and cortisol in blood samples, with DOM animals showed in black and SUBs in red. **(A)** A significant main effect of seasonality was detected on E2 levels (F_(1, 24)_=65.513, p=<0.001, η_p_^2^= 0.732) with highest levels in the birth season. A significant birth rank x seasonality effect emerged on E2 after controlling for lactation during anovulatory season (F_(1, 23)_=3.525, p=0.038, η_p_^2^= 0.133), with SUB females showing higher E2 levels in anovulatory season than DOM animals. **(B)** No significant seasonality, social rank or seasonality x rank effects were detected for cortisol. Graphs show both group (DOM, SUB) mean±SEM (bold lines) and individual (dashed lines) seasonal levels.

There were no significant effects of birth rank on cortisol levels (F_(1, 25)_=0.621, p=0.438, η_p_^2^= 0.024) (Figure 3B). The addition of diet and menarche and the rest of covariates to the model did not change the results. Interestingly, E2 and cortisol levels were positively correlated during birth season (r= 0.391, p= 0.044), despite no associations found during anovulatory (r= 0.287, p= 0.147) or mating (r= 0.316, p= 0.116) seasons.

### Maternal Care: First Month Postpartum

Spearman’s Rank correlations showed significant associations between relative birth rank and frequency of eye-gazing (r=0.714, p=0.006), rejection (r=-0.678, p=0.011), and lipsmacks (r=0.555, p=0.049) with female infants and duration of proximity (r=0.675, p=0.032) and frequency of lipsmacks (r=0.837, p=0.003) with male infants (Figures 4A-4H). Significant associations were also detected between reproductive success rate and frequency of eye-gazing (r=-0.628, p=0.022) with female infants and duration of carry ventral (r=-0.725, p=0.018) with male infants (Figures 5A-5D). Additional associations were found between infant survival rate and proximity ending leave beyond with male infants (r=-0.669, p=0.034) (Figures 5E-5F), as well as between dam’s age at first birth and frequency of retrieve/restrain (r=0.693, p=0.026) with male infants.

**Figure 4.**
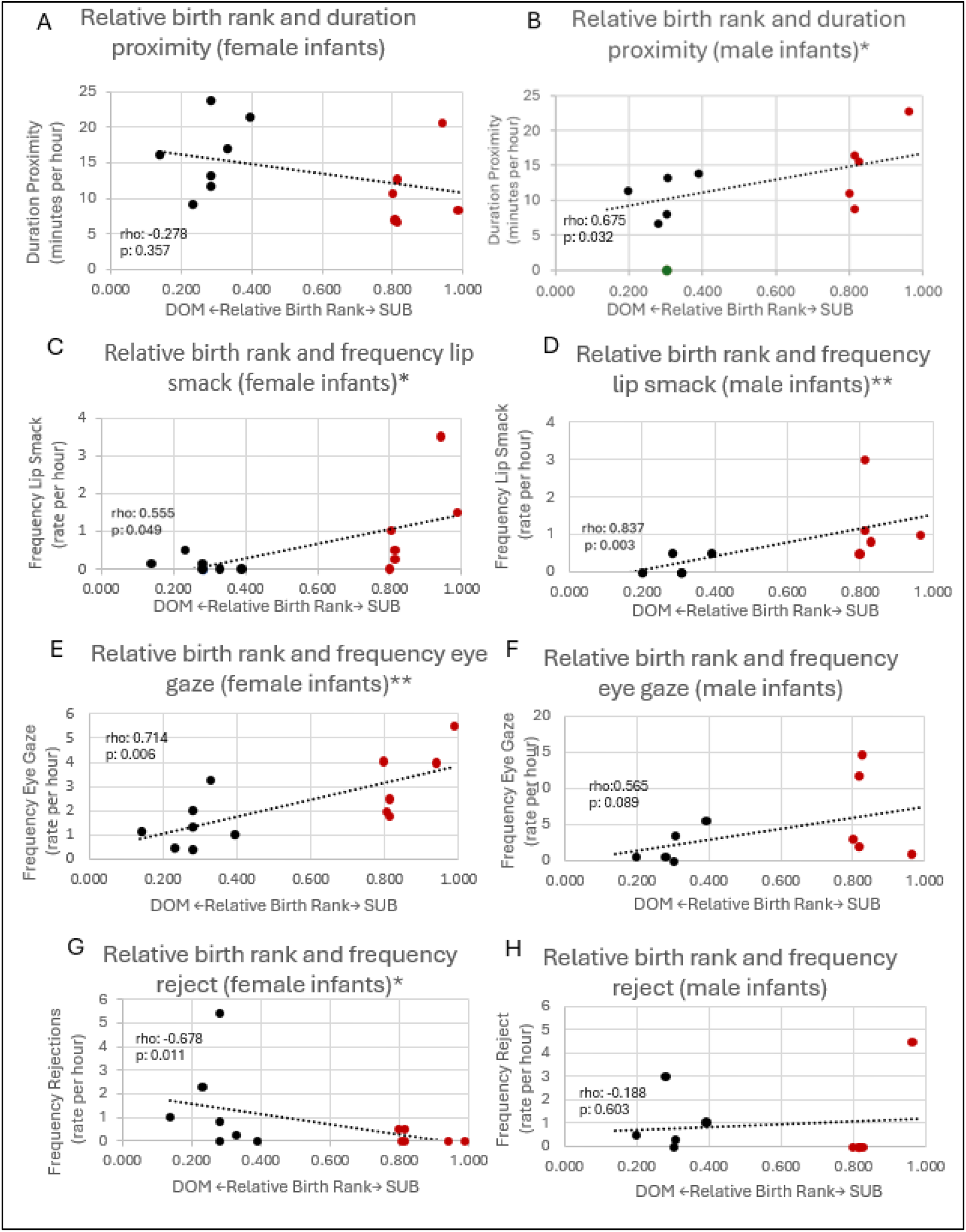
Regression plots showing associations between relative birth rank and mother-infant behaviors. Black data points represent DOM animals; red data points represent SUB individuals. *: p < 0.05; **: p < 0.001.

**Figure 5.**
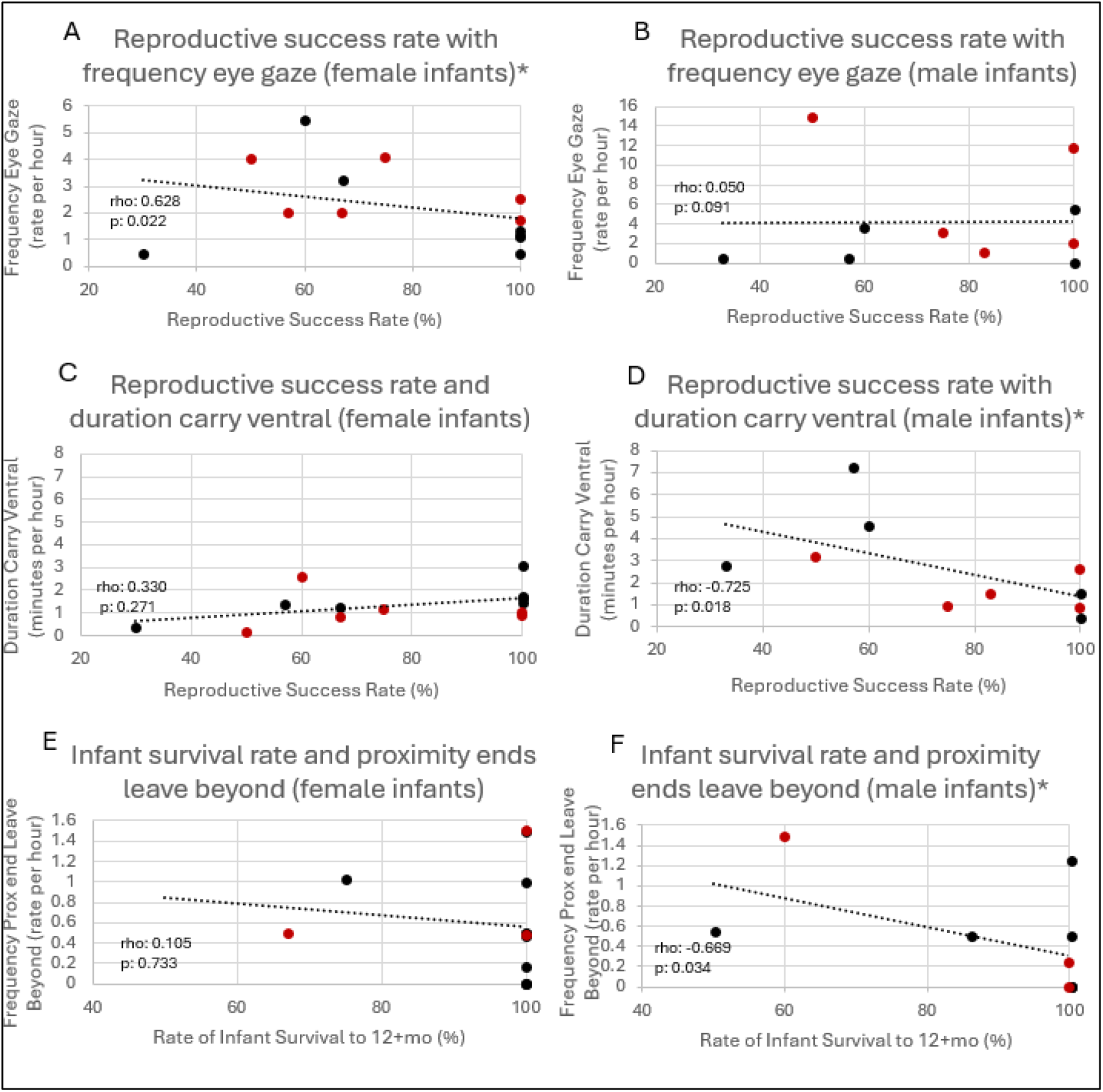
Regression plots showing associations between reproductive success rate, infant survival rate, and mother-infant behaviors. Black data points indicate DOM individuals; red data points indicate SUB individuals. *: p < 0.05; * p < 0.001.

## Discussion

The goal of this study was to investigate long-term effects of chronic social subordination on reproductive function, physiology and maternal care in socially housed female rhesus macaques. In addition to differences in reproductive success and early maternal care between high and low social rank adult females, two main questions in this study involved the underlying biological mechanisms. Thus, we examined (1) whether differences in seasonal levels of E2 related to social rank were associated with reproductive success of female rhesus monkeys and (2) associations between seasonal levels of activity of the HPA and HPG axes by measuring baseline blood levels of E2 and cortisol during the anovulatory, mating and birthing seasons. Our findings suggest long-term effects of social status on reproductive success of adult female rhesus and the likelihood of infant survival. Thus, subordinate females showed lower reproductive success, including less live births and increased rates of stillbirths and of lactation problems postpartum, than dominant females. We then examined seasonal gonadal and stress hormone levels and maternal care as potential predictors behind these social status differences. Subordinate females showed blunted seasonal changes of E2, driven by higher levels in the anovulatory season than dominant animals. Interestingly, higher levels of reproductive hormones (E2) during the anovulatory season predicted lower reproductive success, although the effect was driven by animals lactating at that point. Overall, these findings suggest that female macaques with lower social status show a phenotype consistent with impaired reproductive function, though the biological mechanisms involved need to be further explored. Subordinate females also showed more attentive maternal care of offspring early in life than dominant animals, including higher proximity rates and interactions with infants, likely driven by increased need to protect their infants. Interestingly, a small subset of adult females changed social status during young adulthood (SUB to DOM), which provided the opportunity to test whether this rank switch rescued their reproductive phenotypes. Our findings suggest that even the females that became DOM after a lifelong history of social SUB continue to have reproductive issues, suggesting biological embedding/programming in response to early life adversity.

Females across both groups experienced a significant effect of seasonality on estradiol (E2) levels, which increased from the anovulatory period (Summer) to birth season (Spring). These findings are consistent with previous reports in captive socially housed female macaques living outdoors at the ENPRC colony [33], where a seasonal endocrine rhythm was reported independent of pregnancy and lactation, including gonadal hormone suppression during the anovulatory period. The anovulatory (gonadal suppression) period stems from rhesus monkeys’ seasonal breeding patterns, wherein females have evolutionarily adapted to time mating (ovulation) to match birth and lactation with the time of year where food availability is greatest, given the high levels of energetic output required to raise offspring [53, 54]. This seasonal variation also leads to female synchrony in fertility cycles, thus decreasing male-male competition and increasing female mate choice [55, 56]. During anovulatory season, subordinate females exhibited higher E2 levels than dominants, once we controlled for those who were still lactating from the infant born that year. There was no effect of rank during birth season, but there was an effect of pregnancy and specifically, trimester of pregnancy. The significant increase in E2 between anovulatory and birthing season showed by dominant animals was blunted in subordinate females, although this could be confounded by the fact that dominant females were more likely to have already had an infant from the previous year. This suggests that higher E2 levels in SUB during the anovulatory season could be associated with their lower reproductive success. The female rhesus seasonal HPG endocrine rhythm includes gonadal suppression during the summer anovulatory period (e.g. low E2 blood levels) and rises before ovulation in mating season([33]; the timing and number of ovulations depended on prior levels of E2 and other gonadal hormones. Further studies are needed to examine whether (and how) higher E2 levels during anovulatory season in SUB could negatively impact the number/quality of ovulations or successful pregnancies, given prior evidence that low-ranking females exhibit fewer normal luteal function and ovulations during the breeding season [6].

There were no significant effects of seasonality or social rank on baseline blood cortisol levels, even after controlling for the different confounding factors. Cortisol and E2 levels were positively correlated during birth season, despite no associations found during anovulatory or mating seasons. Our results are consistent with the concurrent increases in E2 and cortisol levels during pregnancy and lactation in multiple species and findings in free-ranging female rhesus monkeys living in Cayo Santiago, where cortisol levels -at least in response to stress-were higher when females were lactating than cycling [38]. The authors also found that the cortisol increase from cycling to lactation was higher in low ranking females, and interpreted the finding as adaptive because heightened activity of stress systems during lactation could be related to increased need to protect the offspring in low than high ranking mothers ([57]), which is also consistent with our observations in SUB mothers in our study, who spent more time in closer proximity to their infants than DOM mothers. Analysis of quality of maternal care showed lower levels of permissiveness (attachment/secure base function) and lower levels of irritability in subordinate than dominant mothers. Alterations in mother-infant attachment measures shown by subordinate females suggest that a mix of biological and maternal care factors could underlie the lower reproductive success of subordinate females in comparison to dominants, and their infants’ decreased likelihood of survival to at least one year of age.

This study has several limitations. The first one is the sample size which, although big for a study with socially housed macaques, limits the statistical power for the analyses of important interactions between factors. It would have also been important to examine the effects of heritable/prenatal factors from the biological mother (DOM, SUB) and of cross-fostering on the outcomes, but we were underpowered to do so. The sole use of females was another limitation of this study; future studies should include both males and females to assess whether social subordination have different effects in male versus female reproductive function. And finally, we could have better captured anovulatory versus ovulatory states of each female if we would have tracked their monthly ovarian cycles through each season (e.g. through monthly measures of E2 and P). Despite these limitations, the findings of the study are interesting and translational.

Improvement in our understanding of social determinants of physical, mental and social health and the risks related to social adversity is critically needed, especially for their consequences on healthy reproductive outcomes that impact future generations.

## Conclusion

Our findings show significantly lower reproductive success rates (number of live births based on access to males; lower infant survival rates) in subordinate than dominant females. Subordinate females also show blunted seasonal changes of the gonadal hormone estradiol (E2), driven by higher E2 levels in the anovulatory season than dominant animals. Subordinate macaques also showed more attentive maternal care of offspring early in life than dominant animals, including higher proximity rates and interactions with infants. Interestingly, higher levels of reproductive hormones (E2) during the anovulatory season was associated with lower reproductive success, although the effect was driven by animals lactating at that point. Overall, these findings suggest that female macaques with lower social status show a phenotype consistent with impaired reproductive function, though the biological mechanisms involved need to be further explored.

## Author Contributions

Conceptualization: M.M. Sanchez, J. Raper, K. Bailey, M.E. Wilson

Methodology: K. Bailey, T. Jonesteller, J. Raper, M.M. Sanchez Analyses: K. Bailey, T. Jonesteller, M.M. Sanchez, Z.A. Kovacs-Balint

Data Curation and Visualization: K. Bailey, T. Jonesteller, Z.A. Kovacs-Balint

Writing-Original Draft Preparation: K. Bailey, M.M. Sanchez

Writing-Review and Editing: K. Bailey, M.M. Sanchez, T. Jonesteller, J. Raper, J. Bachevalier, M.C. Alvarado, Z.A. Kovacs-Balint, M.E. Wilson

Supervision: M.M. Sanchez

Funding Acquisition: M.M. Sanchez, J.R. Raper, M. C. Alvarado, M.E. Wilson All authors have read and agreed to the published version of the manuscript.

## Ethics Statement

All procedures were performed in accordance with the Animal Welfare Act and the U.S. Department of Health and Human Services “Guide for the Care and Use of Laboratory Animals” and the guidelines of the Declaration of Helsinki. The Emory National Primate Research Center is fully accredited by AAALAC International. All studies approved by the Emory Institutional Animal Care and Use Committee (IACUC) under the following protocols: YER-2002302-040516GA, approved on 04/15/2013; YER-2003417-032116GA approved on 3/21/2016; YER-2003417-032317A, approved on 3/23/2017; PROTO202000146, approved on 11/04/2020 and renewed on 10/19/2023.

This article is a revised and expanded version of a poster entitled “Effects of social status on stress physiology and reproductive success in captive female rhesus macaque (*Macaca mulatta*) living in social breeding groups”, which was presented at the annual meeting of the American Society of Primatologists, Riviera Maya, Mexico, September 2024.

## ACKNOWLEDGEMENTS

This study was conducted with invaluable help from Aaron Grey, Ade Falade, Brittney Dockery, Jennifer Whitley, Nathan McDonald and additional research, animal care, colony management and veterinary staff at the Emory National Primate Research Center (ENPRC) Field Station. This project was funded by NIH grants AG070704, HD077623, HD079969 and the NIH’s Office of the Director, Office of Research Infrastructure Programs P51OD011132 (ENPRC Base Grant).

## Data Availability Statement

“Dataset available on request from the authors”

## References

1. Berga, S.L. and T.L. Loucks, The diagnosis and treatment of stress-induced anovulation. Minerva Ginecol, 2005. 57(1): p. 45–54.

2. Warren, M.P. and J.L. Fried, Hypothalamic amenorrhea. The effects of environmental stresses on the reproductive system: a central effect of the central nervous system. Endocrinol Metab Clin North Am, 2001. 30(3): p. 611–29.

3. Baker, S.L., et al., Behavioral and physiological effects of chronic mild stress in female rats. Physiol Behav, 2006. 87(2): p. 314–22.

4. Bethea, C.L., Centeno, M.L., Cameron, J.L., Neurobiology of stress-induced reproductive dysfunction in female macaques. Molecular Neurobiology, 2008. 38: p. 199–230.

5. Kaplan, J.R., et al., Impairment of ovarian function and associated health-related abnormalities are attributable to low social status in premenopausal monkeys and not mitigated by a highisoflavone soy diet. Hum Reprod, 2010. 25(12): p. 3083–94.

6. Pope, N.S., Gordon Thomas P., Wilson Mark E., Age, social rank, and lactational status influence ovulatory patterns in seasonally breeding rhesus monkeys. Biology of Reproduction, 1986. 35(2): p. 353–359.

7. Wagenmaker, E.R., et al., Psychosocial stress inhibits amplitude of gonadotropin-releasing hormone pulses independent of cortisol action on the type II glucocorticoid receptor. Endocrinology, 2009. 150(2): p. 762–9.

8. Xiao, E., L. Xia-Zhang, and M. Ferin, Inadequate luteal function is the initial clinical cyclic defect in a 12-day stress model that includes a psychogenic component in the Rhesus monkey. J Clin Endocrinol Metab, 2002. 87(5): p. 2232–7.

9. Michopoulos, V., et al., Social subordination produces distinct stress-related phenotypes in female rhesus monkeys. Psychoneuroendocrinology, 2012. 37(7): p. 1071–85.

10. Shively, C.A., K. Laber-Laird, and R.F. Anton, Behavior and physiology of social stress and depression in female cynomolgus monkeys. Biol Psychiatry, 1997. 41(8): p. 871–82.

11. Whirledge, S. and J.A. Cidlowski, A role for glucocorticoids in stress-impaired reproduction: beyond the hypothalamus and pituitary. Endocrinology, 2013. 154(12): p. 4450–68.

12. Wilson, M.E., et al., Gonadal steroid modulation of the limbic-hypothalamic-pituitary-adrenal (LHPA) axis is influenced by social status in female rhesus monkeys. Endocrine, 2005. 26(2): p. 89–97.

13. Rivier, C. and S. Rivest, Effect of stress on the activity of the hypothalamic-pituitary-gonadal axis: peripheral and central mechanisms. Biol Reprod, 1991. 45(4): p. 523–32.

14. Avorgbedor, F., et al., Life stressors, hypertensive disorders of pregnancy, and gestational diabetes by race/ethnicity. PLoS One, 2025. 20(4): p. e0321615.

15. Michopoulos, V., et al., Psychophysiology and posttraumatic stress disorder symptom profile in pregnant African-American women with trauma exposure. Arch Womens Ment Health, 2015. 18(4): p. 639–48.

16. Bernstein, I.S., Dominance, aggression and reproduction in primate societies. Journal of Theoretical Biology, 1976. 60(2): p. 459–472.

17. Shively, C.A. and S.M. Day Social inequalities in health in nonhuman primates. Neurobiology of stress, 2015. 1, 156–163 DOI: 10.1016/j.ynstr.2014.11.005.

18. Wilson, M.E., An Introduction to the Female Macaque Model of Social Subordination Stress, in Social inequalities in health in nonhuman primates the biology of the gradient, M.E. Wilson and C.A. Shively, Editors. 2016, Springer International Publishing.

19. Bernstein, I.S. and T.P. Gordon, The function of aggression in primate societies. Am Sci, 1974. 62(3): p. 304–11.

20. Bernstein, I.S., T.P. Gordon, and R.M. Rose, Aggression and social controls in rhesus monkey (Macaca mulatta) groups revealed in group formation studies. Folia Primatol (Basel), 1974. 21(2): p. 81–107.

21. Michopoulos, V., et al., Social subordination impairs hypothalamic-pituitary-adrenal function in female rhesus monkeys. Horm Behav, 2012. 62(4): p. 389–99.

22. Shively, C.A., et al., Social stress-associated depression in adult female cynomolgus monkeys (Macaca fascicularis). Biol Psychol, 2005. 69(1): p. 67–84.

23. Spencer-Booth, Y., The behaviour of group companions towards rhesus monkey infants. Anim Behav, 1968. 16(4): p. 541–57.

24. Howell, B.R., et al., Early adverse experience increases emotional reactivity in juvenile rhesus macaques: relation to amygdala volume. Dev Psychobiol, 2014. 56(8): p. 1735–46.

25. Casabiell, X., et al., Presence of leptin in colostrum and/or breast milk from lactating mothers: a potential role in the regulation of neonatal food intake. J Clin Endocrinol Metab, 1997. 82(12): p. 4270–3.

26. Howell, B.R., et al., Brain white matter microstructure alterations in adolescent rhesus monkeys exposed to early life stress: associations with high cortisol during infancy. Biol Mood Anxiety Disord, 2013. 3(1): p. 21.

27. Dixson, A.F. and C.M. Nevison, The socioendocrinology of adolescent development in male rhesus monkeys (Macaca mulatta). Horm Behav, 1997. 31(2): p. 126–35.

28. Walker, M.L., T.P. Gordon, and M.E. Wilson, Menstrual cycle characteristics of seasonally breeding rhesus monkeys. Biol Reprod, 1983. 29(4): p. 841–8.

29. Du, Y., et al., Seasonal changes in the reproductive physiology of female rhesus macaques (Macaca mulatta). J Am Assoc Lab Anim Sci, 2010. 49(3): p. 289–93.

30. Hinde, K., Richer milk for sons but more milk for daughters: Sex-biased investment during lactation varies with maternal life history in rhesus macaques. Am J Hum Biol, 2009. 21(4): p. 512–9.

31. Wolfensohn, S.a.H. P., Handbook of primate husbandry and welfare. 1st ed. 2005: Wiley-Blackwell.

32. Lee, D.S., A.V. Ruiz-Lambides, and J.P. Higham, Higher offspring mortality with short interbirth intervals in free-ranging rhesus macaques. Proc Natl Acad Sci U S A, 2019. 116(13): p. 6057–6062.

33. Walker, M.L., M.E. Wilson, and T.P. Gordon, Endocrine control of the seasonal occurrence of ovulation in rhesus monkeys housed outdoors. Endocrinology, 1984. 114(4): p. 1074–81.

34. Wilson, M.E., Social dominance and female reproductive behaviour in rhesus monkeys (Macaca mulatta). Animal Behaviour, 1981. 29(2): p. 472–482.

35. Cameron, J.L., Stress and behaviorally induced reproductive dysfunction in primates. Semin Reprod Endocrinol, 1997. 15(1): p. 37–45.

36. Gust, D.A., et al., Relationship between social factors and pituitary-adrenocortical activity in female rhesus monkeys (Macaca mulatta). Horm Behav, 1993. 27(3): p. 318–31.

37. Howell, B.R., et al., Social subordination stress and serotonin transporter polymorphisms: associations with brain white matter tract integrity and behavior in juvenile female macaques. Cereb Cortex, 2014. 24(12): p. 3334–49.

38. Hoffman, C.L., et al., Effects of reproductive condition and dominance rank on cortisol responsiveness to stress in free-ranging female rhesus macaques. Am J Primatol, 2010. 72(7): p. 559–65.

39. McCormack, K.M., et al., The developmental consequences of early adverse care on infant macaques: A cross-fostering study. Psychoneuroendocrinology, 2022. 146: p. 105947.

40. Liu, B.J., et al., Effects of group size and rank on mother-infant relationships and reproductive success in rhesus macaques (Macaca mulatta). Am J Primatol, 2018. 80(7): p. e22881.

41. Hinde, R.A. and Spencerb, Y, Towards Understanding Individual Differences in Rhesus Mother-Infant Interaction. Animal Behaviour, 1971. 19(Feb): p. 165-&.

42. Aarnoudse-Moens, C.S.H., et al., Meta-Analysis of Neurobehavioral Outcomes in Very Preterm and/or Very Low Birth Weight Children. Pediatrics, 2009. 124(2): p. 717–728.

43. Janthakhin, Y., et al., Maternal high-fat diet leads to hippocampal and amygdala dendritic remodeling in adult male offspring. Psychoneuroendocrinology, 2017. 83: p. 49–57.

44. Sullivan, E.L., et al., Chronic Consumption of a High-Fat Diet during Pregnancy Causes Perturbations in the Serotonergic System and Increased Anxiety-Like Behavior in Nonhuman Primate Offspring. The Journal of Neuroscience, 2010. 30(10): p. 3826.

45. Michopoulos, V., D. Toufexis, and M.E. Wilson, Social stress interacts with diet history to promote emotional feeding in females. Psychoneuroendocrinology, 2012. 37(9): p. 1479–1490.

46. Wilson, M.E., et al., Quantifying food intake in socially housed monkeys: Social status effects on caloric consumption. Physiology & Behavior, 2008. 94(4): p. 586–594.

47. Sanchez, M.M., et al., Alterations in diurnal cortisol rhythm and acoustic startle response in nonhuman primates with adverse rearing. Biol Psychiatry, 2005. 57(4): p. 373–81.

48. McCormack, K., et al., Long-Term Effects of Adverse Maternal Care on Hypothalamic-Pituitary-Adrenal (HPA) Axis Function of Juvenile and Adolescent Macaques. Biology (Basel), 2025. 14(2).

49. Raper, J., et al., Neonatal amygdala lesions lead to increased activity of brain CRF systems and hypothalamic-pituitary-adrenal axis of juvenile rhesus monkeys. J Neurosci, 2014. 34(34): p. 11452–60.

50. Wilson, M.E., et al., Social and emotional predictors of the tempo of puberty in female rhesus monkeys. Psychoneuroendocrinology, 2013. 38(1): p. 67–83.

51. Bishop, C.V., et al., Chronically elevated androgen and/or consumption of a Western-style diet impairs oocyte quality and granulosa cell function in the nonhuman primate periovulatory follicle. J Assist Reprod Genet, 2019. 36(7): p. 1497–1511.

52. Altmann, S.A., A field study of the sociobiology of rhesus monkeys, Macaca mulatta. Ann N Y Acad Sci, 1962. 102: p. 338–435.

53. Fooden, J., Systematic Review of the Rhesus Macaque, Macaca Mulatta (Zimmerman, 1780). 2000, Chicago, IL: Field Museum of Natural History.

54. Brockman, D., Seasonality in primate ecology, reproduction, and life-history: an overview, in Seasonality in Primates: Studies of Living and Extinct Human and Non-Human Primates, D. Brockman, Schaik, C, Editor. 2005, Cambridge University Press: New York, NY. p. 3–21.

55. Gogarten, J.F., Koenig, A., Reproductive seasonality is a poor predictor of receptive synchrony and male reproductive skew among nonhuman primates. Behav Ecol Sociobiol, 2013. 67: p. 123–134.

56. Ostner, J., C.L. Nunn, and O. Schulke, Female reproductive synchrony predicts skewed paternity across primates. Behav Ecol, 2008. 19(6): p. 1150–1158.

57. Maestripieri, D., Assessment of danger to themselves and their infants by rhesus macaque (Macaca mulatta) mothers. J Comp Psychol, 1995. 109(4): p. 416–420.

